# “Deciphering the amyloid foldome of TDP-43”

**DOI:** 10.1101/723817

**Authors:** Miguel Mompeán, Emanuele Buratti, Douglas V. Laurents

## Abstract

TDP-43 is an essential regulator of RNA splicing and metabolism and its aggregates play key roles in devastating diseases, including Amyotrophic Lateral Sclerosis (ALS)^1^, Frontotemporal Dementia (FTD) and Limbic-Predominant Age-Related TDP-43 Encephalopathy (LATE)^2^. Besides this pathological aggregation, TDP-43’s oligomerization also serves vital functions^3^, which adds urgency to determine pathological conformations of TDP-43. The recently published cryo-EM study by Cao, Eisenberg and coworkers now reveals amyloid structures of putative pathological aggregates from TDP-43’s C-terminal region^4^. Whereas the Cao *et al*.’s cryo-EM structures contain both the hydrophobic and Q/N-rich segments, the data were interpreted mainly through the lens of hydrophobic contacts. However, the Q/N-rich region can form amyloid on its own^5,6^ and therefore additional considerations of the Q/N-rich segment’s contributions will advance our understanding of TDP-43 aggregation.

---

TDP-43 contains three well-folded domains (residues 1-78, 104-176, and 192-262), and an intrinsically disordered, C-terminal region (residues 265 - 414) (Fig. 1a) containing two evolutionarily conserved segments, suggesting functional importance. These are a stretch of hydrophobic residues (321-340, colored **green** in the sequence shown below), followed by a Q/N-rich, prion-like segment (341-363, in **red**) punctuated by residues with a high turn-forming propensity (<u>underscored</u>):

**Fig. 1.**
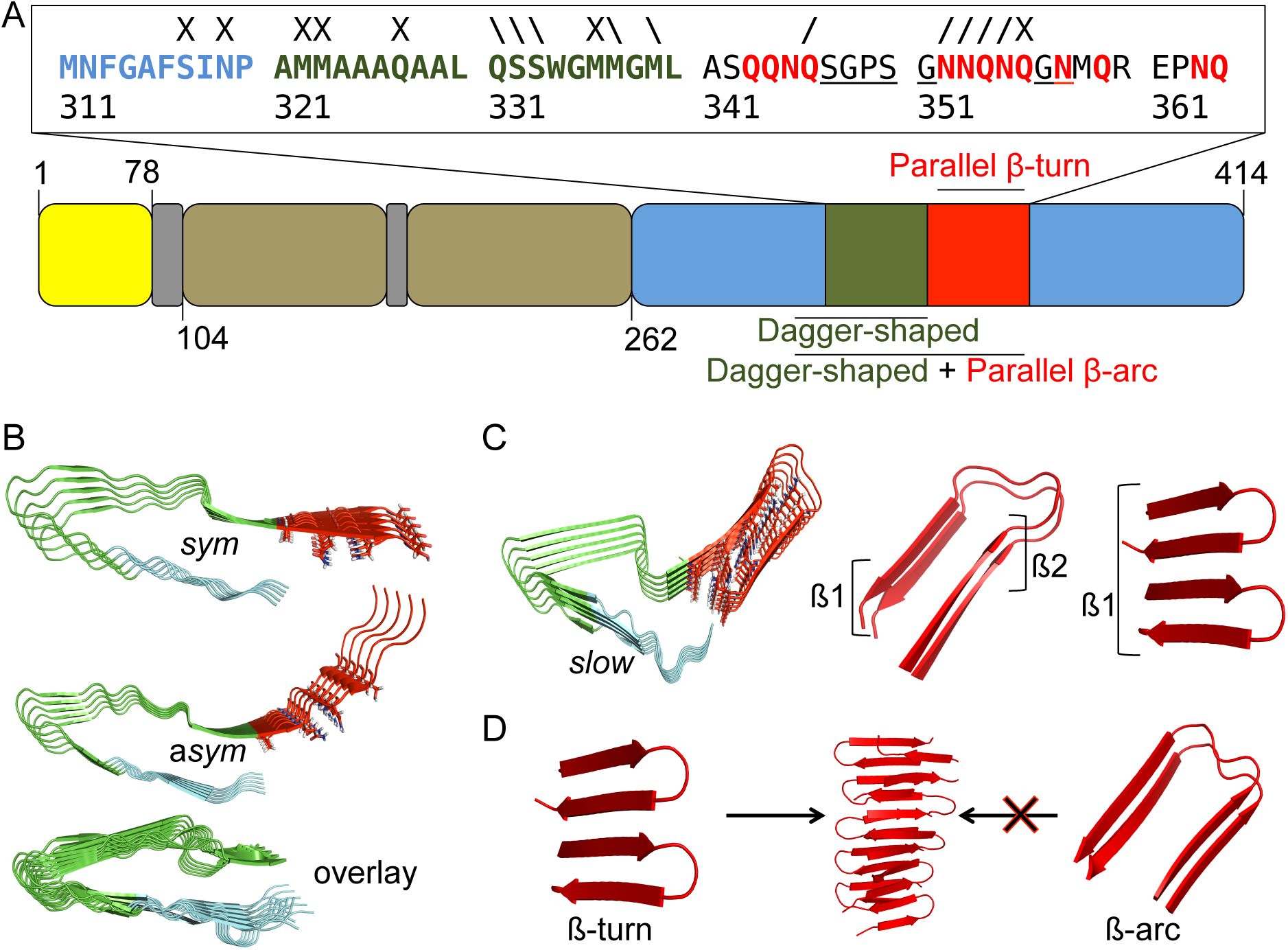
Q/N-rich and hydrophobic segments in TDP-43 aggregation. **a**, Structural domains of TDP-43: N-terminal domain (NTD, in yellow); RNA-recognition motifs (RRMs, in brown); C-terminal domain (in blue), which contains the hydrophobic (green) and Q/N-rich (red) regions that form amyloids with the topologies indicated. Linker regions are shown in gray. The region 311-360 is expanded, as the SegA construct contains these residues, with the pattern of oxidations (Met), phosphorylations (Ser) and deamidations (Gln & Asn) seen by Hasegawa and co-workers^7^ indicated as follows: / = modified in brain 1, \ = modified in brain 2, X = modified in both brains. **b**, Residues within the hydrophobic stretch (321-340 in green) adopt an essentially identical “dagger-shaped” fold in SegA-sym (top), SegA-asym (middle). Residues within the Q/N-rich region (in red) adopt different conformations. The bottom figure illustrates the overlap between SegA-*sym*, SegA-*asym* and SegA-*slow*. **c**, Left: the dagger-shaped fold in SegA-slow is preserved (green), and the Q/N-rich region (red) adopts a hairpin conformation with a β-arc configuration. Middle: each β-strand of the β-arc hairpin contributes to a different β-sheet. Right: the Q/N-rich region adopts a hairpin conformation with a β-turn configuration, where the two β-strands contribute to the same β-sheet**. d**, Hairpins with the β-turn configuration can assemble amyloids (left path), whereas β-arc can not (right path).

~~~
MNFGAFSINP **AMMAAAQAAL QSSWGMMGML** AS**QQNQ**<u>SGPS</u> <u>G</u>**NNQNQ**<u>G</u>**<u>N</u>**M**Q**R EP**NQ**
311 321 331 341 351 361
~~~

Cao *et al.* present the first cryo-EM structures of three polymorphic amyloid fibrils formed by this conserved region of TDP-43, which they called “SegA”. In all three polymorphs (named *sym, asym*, and *slow*), residues within the hydrophobic stretch (321-340) adopt an essentially identical “dagger-shaped” fold (Fig. 1b). Conversely, residues within the contiguous Q/N-rich region (341-360) adopt a variety of conformations, with residues 341-347 being solvent exposed in the two polymorphic structures regarded as “pathologically relevant” (SegA-*sym* and SegA-*asym*) (Fig. 1b). This contrasts previous results by Hasegawa and coworkers, who characterized TDP-43 fibrils derived from two post mortem ALS patient brains using mass spectrometry^7^, and found that Q and N residues within the Q/N-rich stretch, and especially within the 341-348 stretch, were scarcely modified in both brain samples (Fig. 1a). This points to these residues being protected inside the fibrillar core in pathologically relevant, *in vivo* TDP-43 aggregates, which is consistent with our structural model of the TDP-43 amyloid, formed by β-hairpins composed of residues 341-357 assembling with a β-turn configuration^5,6^.

Cao *et al*. also reported the structure of a third polymorph, SegA-*slow*. Unlike SegA-*sym* and SegA-*asym*, the entire Q/N-rich segment is visible in SegA-*slow*, and therefore it could have been compared to the structural model for the Q/N-rich region advanced by Mompeán *et al*.^6^. In particular, the Q/N-rich segment in SegA-*slow* adopts β-hairpin conformation, where residues 341-348 are buried in the fibril core (Fig. 1c), in agreement with the data from brain-derived pathological aggregates discussed above^7^, and contrasting the conformations in SegA-*sym* and SegA-*asym*.

The β-hairpin in the SegA-*slow* structural model has a β-arc configuration, where each of the two β-strands contributes to two different β-sheets (Fig. 1c). By contrast, in our model, β-hairpins are in the β-turn configuration, and the two β-strands contribute to the same β-sheet (Fig. 1c). Interestingly, our previous results indicate that β-arc hairpins formed by the Q/N-rich segment cannot seed the growth of amyloids, whereas the β-hairpin with a β-turn configuration can^6^ (Fig. 1d). Perhaps in the SegA construct, the presence of the 311-320 nonpolar segment (in **blue**, Fig. 1a) confers the flexibility necessary for a preferential hydrophobic-driven assembly (i.e., initial formation of the dagger-shaped structure by residues 321-340), which limits the conformations that the remaining Q/N-rich segment (residues 341-360) may adopt (note that the Q/N-rich region is pruned right at the edge in SegA, Fig. 1a). In this regard, we note that β-arc hairpins are commonly seen in predominantly hydrophobic amyloids, whereas Q-rich fibrils feature hairpins with β-turns^8^. According to the study by Hasegawa and coworkers cited earlier^7^, the M, S and Q+N residues of the hydrophobic stretch that form the dagger-shaped structure in SegA were oxidized, phosphorylated and deamidated, respectively, in both ALS brain samples. This suggests that they are relatively exposed in bona fide TDP-43 fibrils *in vivo* (Fig. 1a), and that Q/N-driven amyloids may be more relevant. There is additional evidence pointing to the ability of the Q/N-rich segment to drive the assembly:

1. TDP-43 constructs with repeats of the Q/N-rich segment are able to induce and reproduce ALS pathological hallmarks in cells^9^.
2. H-bond hypercooperativity among aligned Q/N side chain amide groups are exceptionally strong and decisively stabilize Q/N-rich amyloid structures, including TDP-43, with respect to their hydrophobic counterparts^6,10^.
3. Distinct aggregates of TDP-43 from different pathological tissues are reminiscent of prion strains^11^ In this regard, TDP-43 remains functional when replacing its C-terminal domain with a Q/N-rich prion domain^12^.
4. TDP-43 assembles into a functional amyloid in myogranules^3^, which displays a fibrillar morphology which closely resembles that of the amyloid with the β-turn β-hairpin conformation formed by the Q/N-rich region^6^. Both are recognized by the A11 antibody.

Finally, in addition to forming homo-oligomeric assemblies, TDP-43 interacts with Ataxin-2^13^ and polyQ segments through its C-terminal region^14^, and also recruits Nucleoporins (Nups) into TDP-43+Nups co-aggregates in disease^15^. These interactions among Q/N-rich domains strongly suggest that hybrid amyloids are formed, which could resemble the recently determined structure of the RIPK1-RIPK3 human necrosome, where Q residues are key to RIPK1-RIPK3 assembly^16^. The structures presented here by Cao *et al.* undoubtedly reveal an important portion of the TDP-43 aggregate foldome, highlighting the role of the hydrophobic segment. Nonetheless, based on the considerations mentioned here, the role of the Q/N-rich region in pathologically relevant structures might well be more central.

## Acknowledgements

This work was supported by a Junior Leader Project LCF/BQ/PR19/11700003 from “La Caixa” Foundation (MMG), project PathensTDP-ArisLA from the Italian Research Foundation for ALS (EB), and project SAF2016-76678-C2-2-R from the Spanish Ministry of Economy and Competitivity (DVL).

## Author contributions

M.M., E.B., and D.V.L. edited and co-wrote the manuscript.

## Competing interest

The authors declare no competing interests.

## References

1. Neumann, M. et al. Ubiquitinated TDP-43 in Frontotemporal Lobar Degeneration and Amyotrophic Lateral Sclerosis. Science 314, 130–633 (2006).

2. Nelson, P. T. et al. Limbic-predominant age-related TDP-43 encephalopathy (LATE): consensus working group report. Brain. 146(6), 1503–1627 (2019).

3. Vogler, T. O. et al. TDP-43 and RNA form amyloid-like myo-granules in regenerating muscle. Nature 563, 508–513 (2018).

4. Cao, Q., Boyer, D. R., Sawaya, M. R., Ge, P. & Eisenberg, D. S. Cryo-EM structures of four polymorphic TDP-43 amyloid cores. Nat. Struct. Mol. Biol. 26, 619–627 (2019).

5. Mompeán, M. et al. Structural characterization of the minimal segment of TDP-43 competent for aggregation. Arch. Biochem. Biophys. 545, 53–62 (2014).

6. Mompeán, M. et al. Structural evidence of amyloid fibril formation in the putative aggregation domain of TDP-43. J. Phys. Chem. Lett. 6, 2608–2615 (2015).

7. Kametani, F. et al. Mass spectrometric analysis of accumulated TDP-43 in amyotrophic lateral sclerosis brains. Sci. Rep. 6, 23281 (2016).

8. Hoop, C. D. et al. Huntingtin exon 1 fibrils feature an interdigitated β-hairpin-based polyglutamine core. Proc. Natl. Acad. Sci. 113, 1546–1551 (2016).

9. Budini, M. et al. Cellular model of TAR DNA-binding protein 43 (TDP-43) aggregation based on its C-terminal Gln/Asn-rich region. J. Biol. Chem. 287, 7512–7525 (2012).

10. Mompeán, M., Nogales, A., Ezquerra, T. A., & Laurents, D. V. Complex system assembly underlies a two-tiered model of highly delocalized electrons. J. Phys. Chem. Lett. 6, 1831–1864 (2016)

11. Laferrière, F. et al. TDP-43 extracted from frontotemporal lobar degeneration subject brains displays distinct aggregate assemblies and neurotoxic effects reflecting disease progression rates. Nat. Neurosci. 22, 65 (2019).

12. Wang, I. F. et al. The self-interaction of native TDP-43 C terminus inhibits its degradation and contributes to early proteinopathies. Nat. Comunn. 3, 766 (2012).

13. Becker, L. A. et al. Therapeutic reduction of ataxin-2 extends lifespan and reduces pathology in TDP-43 mice. Nature 544, 367–371 (2017).

14. Fuentealba, R. A. et al. Interaction with polyglutamine aggregates reveals a Q/N-rich domain in TDP-43. J. Biol. Chem. 285, 26304–26314 (2010).

15. Chou, C. C. et al. TDP-43 pathology disrupts nuclear pore complexes and nucleocytoplasmic transport in ALS/FTD. Nat. Neurosci. 21, 228–239 (2018).

16. Mompeán, M. et al. The structure of the necrosome RIPK1-RIPK3 core, a human hetero-amyloid signaling complex. Cell 173, 1244–1253 (2018).

